# Magnetoelectric Microrobots for Spinal Cord Injury Regeneration

**DOI:** 10.1101/2024.08.06.606378

**Authors:** Hao Ye, Jingjing Zang, Jiawei Zhu, Denis von Arx, Vitaly Pustovalov, Minmin Mao, Qiao Tang, Andrea Veciana, Harun Torlakcik, Elric Zhang, Semih Sevim, Roger Sanchis-Gual, Xiang-Zhong Chen, Daniel Ahmed, Maria V. Sanchez-Vives, Josep Puigmartí-Luis, Bradley J. Nelson, Stephan C. F. Neuhauss, Salvador Pané

## Abstract

Regenerative medicine continually seeks effective methods to address spinal cord injuries (SCI), which are known for their limited regenerative potential. Despite advances in neural progenitor cell (NPC) transplants for spinal cord injuries, challenges related to graft survival, reliable *in vivo* differentiation, and neural integration significantly hinder real functional recovery and limit clinical outcomes. This study introduces ‘NPCbots’, biohybrid microrobots engineered by integrating human-induced pluripotent stem cell-derived NPCs with magnetoelectric nanoparticles composed of cobalt ferrite-barium titanate. These enable magnetic navigation and neuronal stimulation, enhancing targeted therapeutic interventions. Our lab-on-a-chip system allows for the mass production of NPCbots, ensuring their differentiation and biocompatibility. Remarkably, in a zebrafish model of SCI, NPCbots stimulated by an alternating magnetic field demonstrated rapid *in vivo* differentiation and integration into damaged neural pathways, significantly enhancing neural regeneration. Within three days, injured zebrafish treated with NPCbots exhibited almost normal swimming behavior and significantly improved exploratory behavior, showcasing the potential of NPCbots to swiftly repair neural structures and restore the central nervous system’s functionality in spinal cord injury models through non-invasive means. Additionally, precise *in vitro* and *in vivo* manipulation of NPCbots indicates their broader application in various neurodegenerative disorders, offering a promising route for effective spinal cord and neurological recovery.

## Introduction

Cell therapies represent a burgeoning segment of biotechnology, aimed at curing diseases through targeted cell delivery to specific tissues. These therapies are diverse, encompassing tissue regeneration, restoration of biological functions, and enhancement of the body’s intrinsic disease-fighting capabilities^1, 2^. Unlike many drugs and biological agents, delivered cells can traverse biological barriers, respond to and adapt to biological signals, and specifically target cells, tissues, or organs^3, 4^. While the safety and efficacy of cell therapies have been enhanced by minimally invasive targeted cell delivery techniques, therapeutic efficacy is still suboptimal, with low cell survival rates (approximately 5%) and unreliable *in vivo* differentiation into targeted cells^5–9^. For instance, selective colonization by neural stem cells (NSCs), which aim to differentiate only upon reaching the target area, addresses these issues^10, 11^. Neurotrophins, such as nerve growth factor (NGF), typically act as chemical stimuli^12^, but are less effective *in vivo* due to rapid diffusion rates, delivery challenges, and short half-lives^13, 14^. These limitations might be overcome by directly stimulating cells to initiate neuronal differentiation at specific post-delivery times^15–17^. Various external stimuli, including light^18^, magnetic fields^19, 20^, and sound waves^21^, have been utilized to induce or enhance targeted neuronal differentiation. Among these, magnetic fields are uniquely capable of deep tissue penetration with minimal attenuation due to the low magnetic susceptibility of biological materials^22–25^.

In recent years, the approach to treating spinal cord injuries (SCI) increasingly focused on the potential benefits of neural progenitor cell (NPC) transplants^26^. Despite considerable progress, obstacles related to graft survival, controlled differentiation, and neural integration continue to severely constrain therapeutic successes and the attainment of real functional recovery^27^. Our study introduces neural progenitor cell-based microrobots (NPCbots) – a novel class of biohybrid microrobots. The unique construction of CoFe_2_O_4_-BaTiO_3_ (CFO-BTO) nanoparticles, through cobalt ferrite and barium titanate synthesis, provides dual functionalities of magnetic manipulation and magnetoelectric stimulation. This enhances both the operational efficiency and therapeutic impact of the microrobots. Techniques such as high-angle annular dark-field scanning transmission electron microscopy (HAADF-STEM) and energy-dispersive X-ray (EDX) spectroscopy mapping, verifying the integrity of the core-shell structure and effective magnetoelectric coupling. These CFO-BTO magnetoelectric core-shell nanoparticles were then integrated with human-induced pluripotent stem cell (iPSC)-derived NPCs, enabling precise *in vivo* magnetic navigation and specific neuronal differentiation within complex biological settings. The fabrication of NPCbots using a bidirectional continuous-flow microfluidic device, which facilitates their mass production (**Fig. 1a**).

**Fig. 1.**
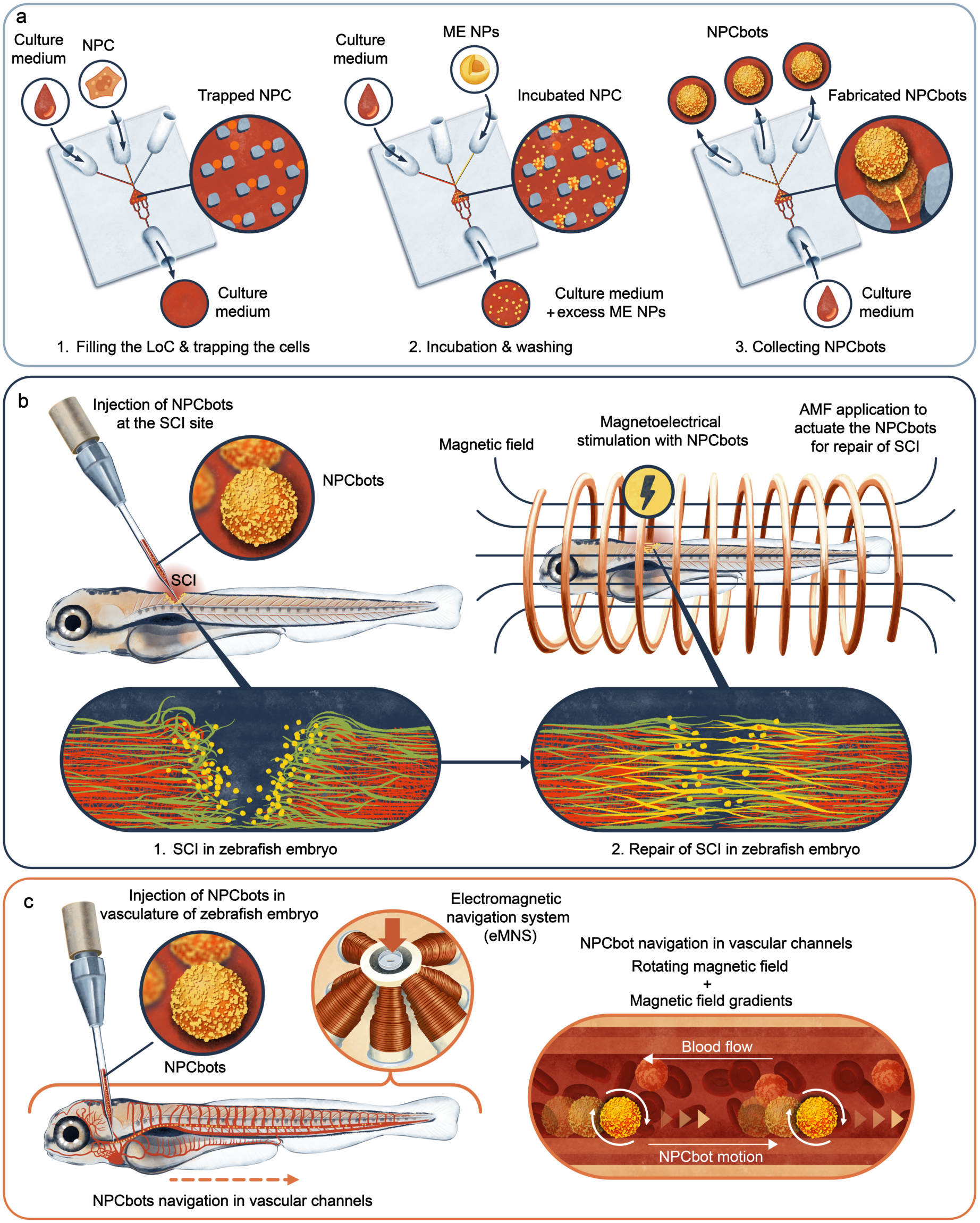
Schematic representation of the neural progenitor cell-based microrobots (NPCbots) fabrication and application processes. **a**, Overview of the NPCbots’ fabrication steps, including neural progenitor cells (NPCs) trapping, incubation of NPCs with cobalt ferrite-barium titanate (CFO-BTO) nanoparticles, collection of the NPCbots. **b**, Application of an alternating magnetic field (AMF) to the zebrafish larvae spinal cord injury model following NPCbot injection. **c**, *In vivo* manipulation of the NPCbot within the dorsal artery (DA) of zebrafish larvae, depicting the NPCbot’s rolling motion through the targeted area within the zebrafish larvae.

Our approach is further illustrated by the deployment of these NPCbots within a zebrafish model of spinal cord injury. By integrating magnetic navigation with *in vivo* neuronal stimulation, we not only enhance regenerative capabilities but also significantly improve functional recovery, as evidenced through comprehensive behavioral and molecular assessments (**Fig. 1b-c**). The use of alternating magnetic fields (AMF) to stimulate the NPCbots results in marked neuronal and astrocytic differentiation, effectively leveraging the magnetoelectric properties of CFO-BTO nanoparticles to direct cellular behavior and promote neural regeneration (**Fig. 1b**). This study contributes to the fields of nanotechnology and regenerative medicine by providing a new approach to the treatment of neurodegenerative diseases. It introduces a platform that allows for targeted and efficient restoration of neural functions through non-invasive means. By utilizing microfluidic technologies – *i.e.*, lab-on-a-chip (LoC) devices – and magnetoelectric nanoparticles, NPCbots represent an advancement in therapeutic applications. They offer promising potential for improving medical treatments for spinal cord injuries and related conditions.

## RESULTS AND DISCUSSION

### Fabrication and Characterization of NPCbots

LoC devices have realized the rapid and efficient mass production of biohybrid microrobots. In this investigation, we detail the fabrication process of these innovative microrobots, as illustrated in **Fig. 1a**. Each microrobot incorporates CFO-BTO and human iPSC-derived NPCs. The CFO-BTO nanoparticles are fabricated by synthesizing a magnetostrictive cobalt ferrite (CFO) core and encasing it in a piezoelectric barium titanate (BTO) shell, where the magnetic field-induced strain in CFO is converted into an electrical output by BTO, thereby enabling magnetoelectric coupling^28^. This configuration is specifically designed to facilitate magnetic manipulation and *in vivo* neuronal magnetoelectric stimulation of the microrobot.

The synthesis of these CFO-BTO core-shell magnetoelectric nanoparticles begins with a co-precipitation process, followed by hydrothermal and sol-gel methods. Initially, cetyltrimethylammonium bromide (CTAB) is dissolved in deionized water, with subsequent additions of ferric chloride hexahydrate and cobalt(II) chloride. The formation of CFO nanoparticles is initiated by adding sodium hydroxide, followed by hydrothermal treatment in a sealed autoclave to enhance crystallinity. The BTO shell is then formed using a sol-gel technique, involving a precursor solution of barium carbonate, citric acid, and titanium isopropoxide. This precursor is mixed with the CFO nanoparticles to form a gel, which is subsequently dried and annealed to complete the synthesis of the CFO-BTO core-shell nanoparticles.

Characterization of these nanoparticles is achieved using HAADF-STEM coupled with EDX mapping. This confirms the core-shell structure, revealing a CFO core rich in Co, Fe, and O, and a BTO shell containing Ba, Ti, and O elements (**Fig. 2a**). The dimensions of the core and shell are 99.4 ± 4.2 nm and 7 ± 3.5 nm, respectively. X-ray diffraction (XRD) analyses further confirm the phase purity and crystalline structure of the nanoparticles, which identified characteristic reflections corresponding to the ferrite CFO phase of the 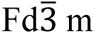 space group (ICDD:98-016-7445) and the ferroelectric BTO phase of the P4mm space group (ICDD:98-001-5453)^29^, with no impurities or intermediate phases detected (**Fig. 2b**). Furthermore, the magnetoelectric coupling properties of the nanoparticles were explored by assessing the local piezoresponse under magnetic fields. Application of a 50 mT DC in-plane magnetic field altered the local piezoelectric hysteresis loops, demonstrating significant magnetoelectric coupling in the CFO–BTO nanoparticles. This was evidenced by the notable shifts in both the positive and negative coercive voltages (dV=1.81V) with respect to no-field, suggesting that the strain generated within the CFO (magnetostrictive phase) effectively transferred to the BTO (piezoelectric phase), thereby facilitating polarization^28, 30^. The observed asymmetry in the piezoresponse loops, evident from an offset under the magnetic influence, likely resulted from an electric field generated through the magnetoelectric effect, validating the strain-mediated magnetoelectric functionality within these core-shell nanoparticles (**Fig. 2c,d**).

**Fig. 2.**
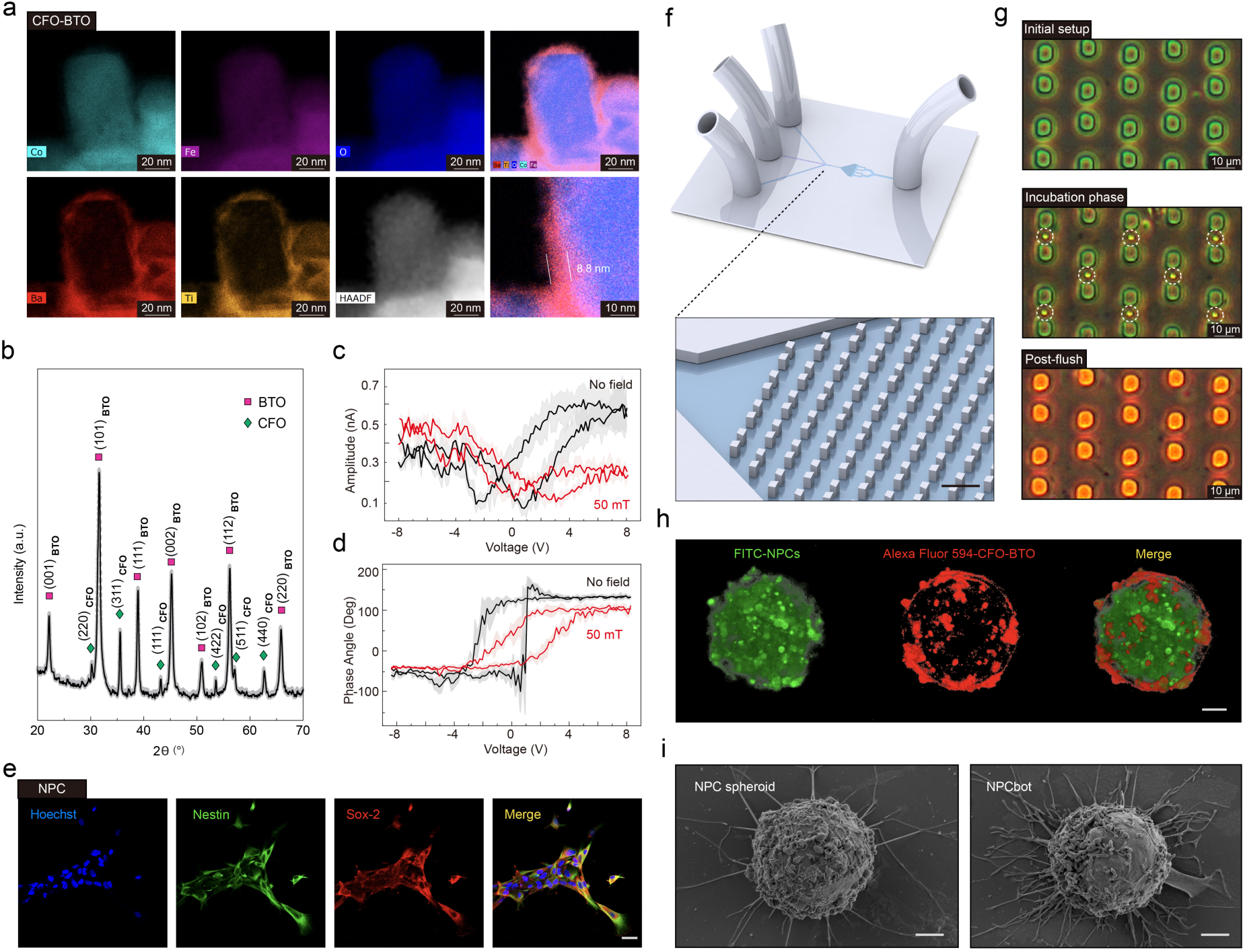
Fabrication and Characterization of NPCbots. **a,** High-angle annular dark-field scanning transmission electron microscopy (HAADF-STEM) and energy-dispersive X-ray (EDX) analyses of CFO-BTO core–shell nanoparticles. **b**, X-ray diffraction (XRD) patterns confirming the crystalline structure of CFO-BTO powders. **c-d,** Magnetic-field-dependent local piezo force microscopy (PFM) hysteresis loops (amplitude and phase in **c** and **d**, respectively) from CFO-BTO core-shell nanoparticles, with (red) and without (black) an applied magnetic field of 50 mT (n = 3). **e**, Immunofluorescence staining for Nestin and Sox-2 in NPCs, highlighting progenitor cell markers (scale bar = 20 μm). **f,** Lab-on-a-chip (LoC) device design featuring a bidirectional continuous-flow microfluidic system that includes a large trapping chamber for cell capture and incubation, three inlets for introducing growth medium, cells, and nanoparticles, and a single outlet which can also function as an inlet during the collection of NPCbots. **g,** Optical microscopy image documenting the fabrication process of NPCbots. **h**, Confocal microscopy images of NPCbots labeled with Alexa Fluor 594-conjugated CFO-BTO nanoparticles and FITC-labeled NPCs after 30 minutes of incubation (scale bar = 2 μm). **i,** Scanning electron microscopy (SEM) images of an NPC spheroid and an assembled NPCbot, demonstrating the structural integrity and surface morphology (scale bar = 2 μm).

The stemness of NPCs is critical for the fabrication of NPCbots. Nestin and Sox-2 are extensively characterized as neural progenitor cell markers and are not present in mature neurons^31^. Both markers colocalize in NPCs, confirming that the neural progenitor cells retain their stem-like properties without undergoing differentiation (**Fig. 2e**).

To construct the NPCbots, a bidirectional continuous-flow microfluidic device was utilized, as depicted in **Fig. 2f** and **Supplementary Fig. 1**. This device is engineered with a large trapping chamber, specifically optimized for capturing and incubating cells to synthesize functional microrobots. It incorporates three inlets on one side for the introduction of growth medium, cells, and nanoparticles, while a single outlet on the opposing side also functions inversely as an inlet during the collection phase of microrobots (**Fig. 1a**). The microfluidic trapping chamber is designed to have a triangular shape for two reasons: it maximizes trapping efficiency based on the probabilistic distribution of cells within the fluid stream and facilitates an accelerated reverse flow during the collection phase, which is critical for preventing clogging at the chamber’s entrance. The traps, with a minimum gap of 6 µm, are finely tuned to ensnare a broad spectrum of cell sizes while minimizing flow resistance that might allow cells to evade capture. The microfluidic chamber is designed with a height of 8-10 µm to prevent the simultaneous trapping of multiple cells at one site. During the NPCbots’ fabrication (**Fig. 1a**), NPCs are ensnared within these traps via a controlled low-flow-rate microfluidic stream. This is followed by the introduction of CFO-BTO flow for *in situ* incubation within the device. After incubation, the outlet of the device is reconfigured to serve as an inlet, using a pump to facilitate a reverse flow that discharges the cultivated microrobots from their traps. The original inlets are then repurposed as outlets for microrobot collection. This innovative bidirectional design not only simplifies the trapping and release of microrobots but also supports their mass production, enhancing the efficiency and convenience of biohybrid microrobot fabrication (**Fig. 2g**).

The viability of the synthesized NPCbots was assessed by introducing varying concentrations of CFO-BTO nanoparticles, selecting a concentration of 33 mg/mL (1 ng CFO-BTO per cell) based on live/dead assay fluorescence images and MTT assays, which demonstrated over 85% viability (**Supplementary Fig. 2**). Further investigations explored the effects of incubation time on cellular uptake of CFO-BTO. To fluorescently label CFO-BTO, nanoparticles were initially coated with (3-Aminopropyl)triethoxysilane (APTES), facilitating amino group attachment to their surfaces. Spectral analysis confirmed the presence of amine groups, as evidenced by the Si-O-Si signal derived from the CFO-BTO-conjugated APTES (**Supplementary Fig. 3**). Subsequently, these APTES-coated nanoparticles were incubated with Alexa Fluor™ 594 NHS Ester (Succinimidyl Ester), which promoted the formation of stable amide bonds between the NHS ester and amine groups. After a 1-hour incubation at 37°C, significant cellular uptake by NPCs was observed, determining a 30-min incubation period as optimal for NPCbot fabrication to ensure maximum interaction with the cell membrane, potentially influencing membrane channels (**Supplementary Fig. 4**).

Following the post-labeling of CFO-BTO with Alexa Fluor 594, there were negligible changes in the size and zeta potential of the nanoparticles (**Supplementary Fig. 5**), facilitating further exploration of their distribution. The NPCbots were subsequently stained using a live/dead staining kit, incorporating a live dye for visualization. Confocal microscopy enhanced by z-stack analysis documented detailed distribution patterns of the labeled CFO-BTO on the NPC surface (**Fig. 2h**). Scanning Electron Microscopy (SEM) was employed to assess the morphological differences between NPCbots and NPC spheroids (without CFO-BTO incubation). Further SEM analysis verified that the morphological integrity of the NPCs was preserved following incubation with CFO-BTO, as indicated by the retention of characteristic cellular microstructures such as filopodia (**Fig. 2i**). These results collectively demonstrate that CFO-BTO nanoparticles can adhere to the surfaces of NPC membranes without compromising cellular architecture, thereby supporting their potential utility in enhancing further differentiation capabilities.

### Stimulation and differentiation of NPCbots via alternating magnetic fields

To investigate the differentiation potential of NPCbots under specific conditions, we employed an AMF stimulation protocol. Our experimental setup involved placing samples within a solenoid coil with an internal diameter of 5 cm, operating in a resonant tank configuration (**Fig. 3a**). This configuration generated a magnetic field with an amplitude of 20 mT and a frequency of 1.18 kHz at the center of the coil. The AMF was applied for two hours, twice daily, for two days to realize the NPCbots differentiation. We used Fibroblast Growth Factor 2 (FGF-2), a neuronal differentiation inducer, as a positive control in these experiments. Immunofluorescence imaging conducted post-AMF treatment revealed a shift of NPCbots towards neuronal cell types. **Fig. 3b-d** showcase these transformations, accompanied by an enhanced expression of neuronal markers, βIII-tubulin and microtubule-associated protein 2 (MAP2). Quantitative analysis indicated a significant increase in the expression of both markers, with mean fluorescence intensity rising by 6.6-fold and 19.1-fold for βIII-tubulin and MAP2, respectively, compared to controls without AMF treatment. These markers are integral to neuronal cell structure and function^32^, signifying effective neural induction through AMF stimulation. Further detailed in **Fig. 3e-g**, NPCbots also demonstrated a significant upregulation of α-tubulin and glial fibrillary acidic protein (GFAP) post-treatment, with mean fluorescence intensity increases of 15.8-fold and 15.7-fold, respectively. α-Tubulin is indicative of neurite outgrowth, while GFAP is a marker for astrocytes^33, 34^, indicating that the NPCbots not only differentiate into neurons but also promote astrocytic lineage development under the influence of AMF. Additionally, significant improvements were also observed in the NPC+CFO-BTO group treated with AMF, mirroring these enhancements. These results elucidate that the magnetoelectric properties of the CFO-BTO nanoparticles, situated on the NPC membrane within the NPCbots, respond to an AMF by generating electric charges, which likely influence calcium channel activity and trigger calcium influx. This influx is presumed to initiate signaling pathways that foster the following differentiation, underscoring the potential of AMF to regulate cellular behavior, which provides a non-invasive method to direct cellular fate.

**Fig. 3.**
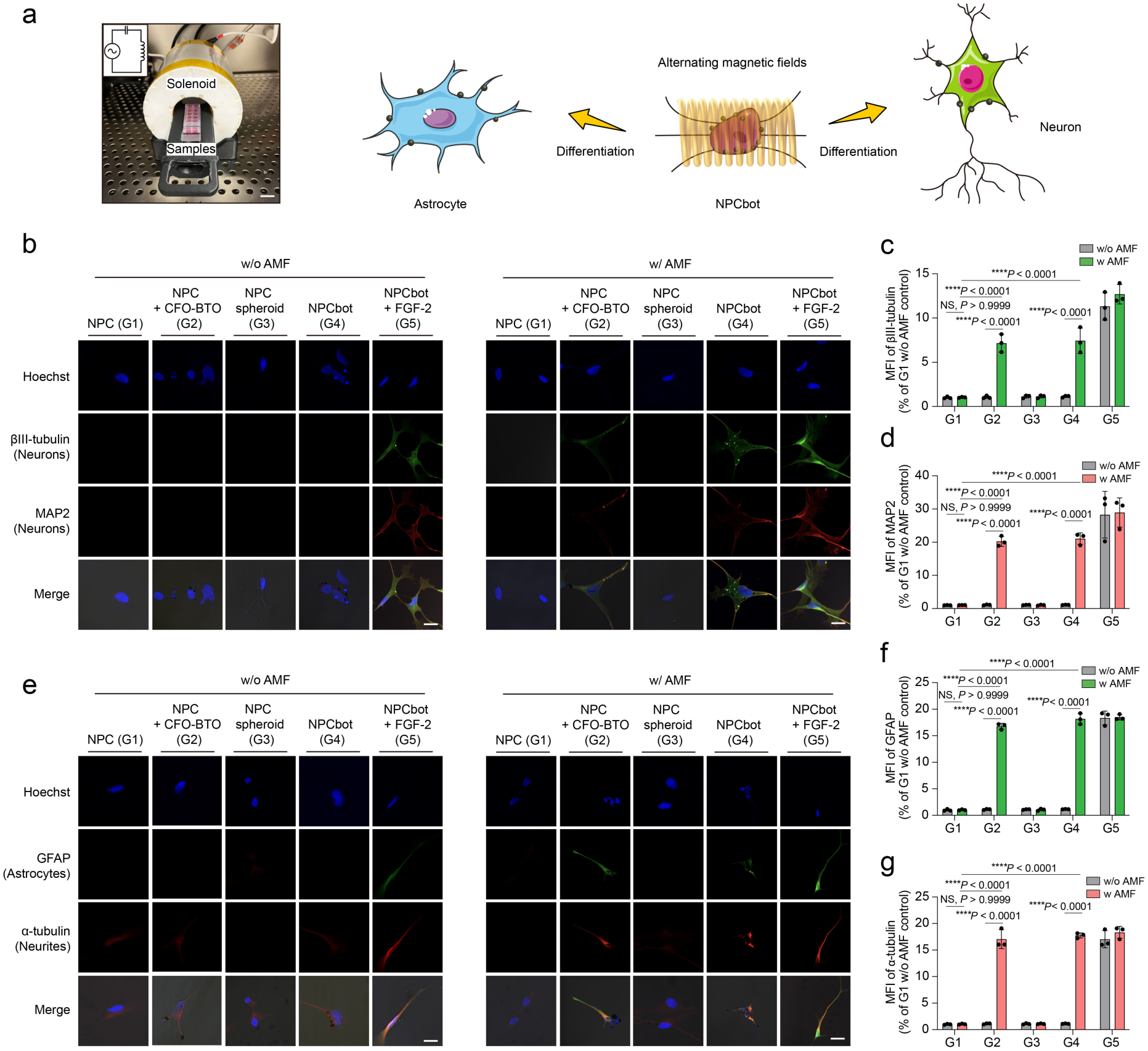
Induction of NPCbots differentiation using alternating magnetic fields. **a**, Schematic representation of the experimental setup for differentiation induction. NPCbots were exposed to alternating magnetic fields generated by a solenoid coil with a 5 cm internal diameter (Scale bar = 20 mm). The coil operated within a resonant tank circuit, producing a magnetic field with a peak amplitude of 20 mT and a frequency of 1.18 kHz at the center of the solenoid. **b-d**, Immunofluorescence analysis of NPCbots showing expression of neuronal markers. Stained for βIII-tubulin and MAP2 to identify differentiated neurons. Quantitative assessment through mean fluorescence intensity (MFI) across all samples (n = 3, means ± s.e.m.). Scale bar = 8 μm. **e-g**, Immunofluorescence characterization of NPCbots stained for α-tubulin and GFAP, markers of differentiated neurites and astrocytes, respectively. MFI was measured for each marker across all samples (n = 3, means ± s.e.m.). Scale bar = 8 μm. Statistical significance was calculated via one-way ANOVA with a Tukey post-hoc test (**Fig. 3c, d, f, g**). **** p< 0.0001 versus control.

### Efficacy of NPCbot Treatment in Zebrafish Spinal Cord Regeneration

To assess the therapeutic efficacy of NPCbot treatment, we established a zebrafish spinal cord injury model. This model was utilized to evaluate the regenerative potential of NPCbots administered via injection and subsequently stimulated with an AMF over a period of five days. As depicted in **Fig. 4a**, the experimental timeline was designed to allow for a comprehensive assessment of recovery following injury at 2 dpf (days postfertilization). Stimulation was conducted for two hours each day from 2 to 4 dpf. The magnetic field direction was oriented parallel to the longitudinal axis of the zebrafish. Employing whole-mount immunohistochemistry, we documented the therapeutic benefits in zebrafish treated with NPCbots (**Fig. 4b**). Notably, group treated with CFO-BTO nanoparticles (G3) exhibited enhanced fluorescence intensity of GFAP at the injury site (**Fig. 4c**), indicative of an active astrocytic response and neural repair. This was coupled with a noticeable reduction in lesion distance (**Fig. 4d**), which may be attributed to the electric charges generated by CFO-BTO under the influence of AMF, which in turn, facilitated the regenerative process. Correspondingly, these findings are consistent with the NPC differentiation results from our *in vitro* cell stimulation test. A marked recovery was most pronounced in the NPCbot-treated group (G5). α-Tubulin staining demonstrated a robust reformation of neuronal structures, crucial for maintaining neural function and integrity. Concurrently, GFAP staining of the NPCbot-treated group (G5) highlighted a significant presence of neural connections compared to the control groups (injured (G2), CFO-BTO treated (G3), and NPC treated (G4)), showcasing the greatest corrected total GFAP fluorescence and the smallest average distance at the injury site. This underscores that combining the NPCbot strategy is crucial in aiding the zebrafish’s spinal cord to repair itself (**Fig. 4b-d**).

**Fig. 4.**
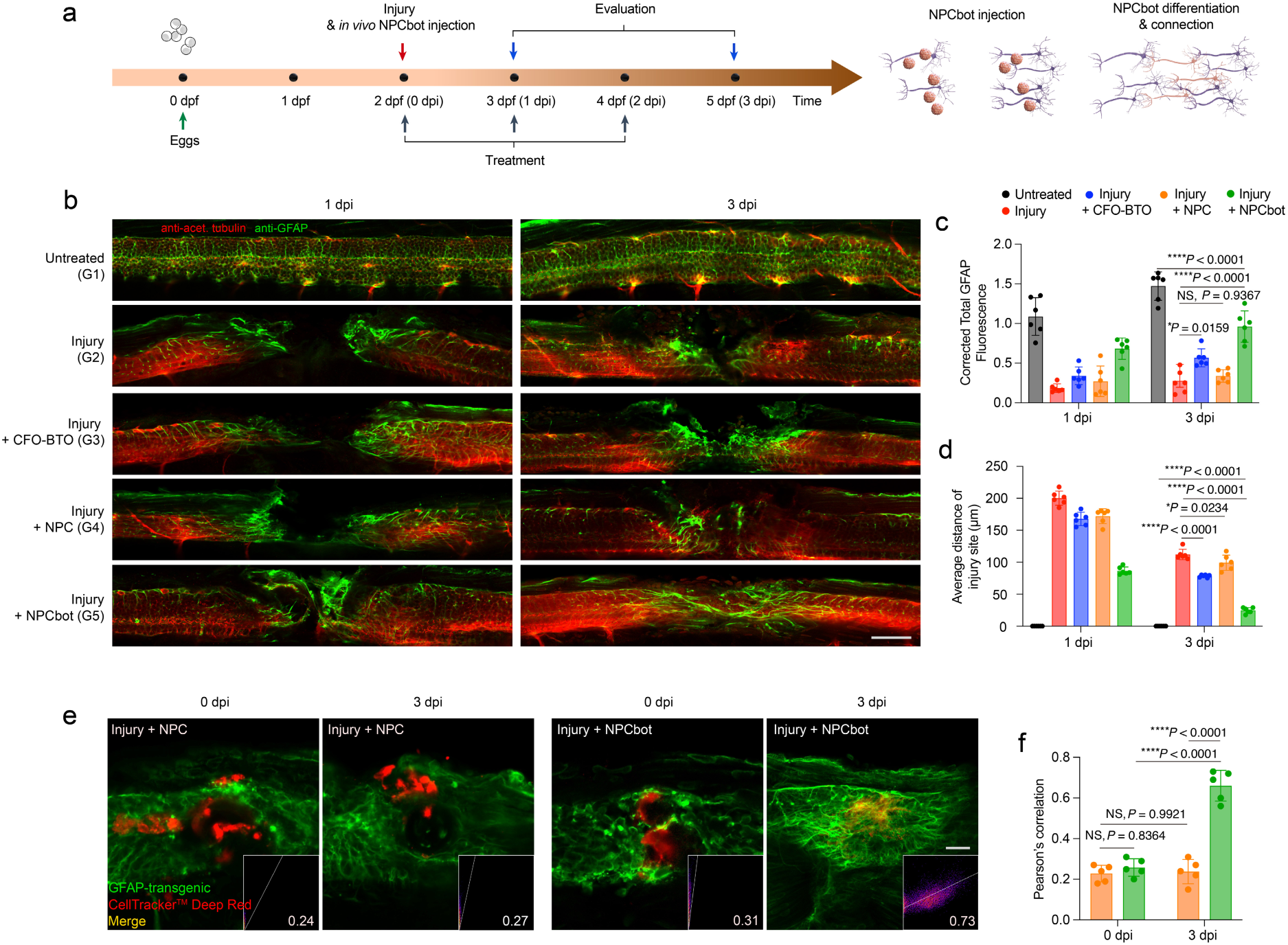
Therapeutic efficacy of NPCbot in treating zebrafish spinal cord injuries. **a**, Schematic representation of the experimental design for treatment processes. **b**, Representative whole-mount immunohistochemistry images demonstrating co-labeling of neurites (anti-acetylated tubulin, red) and the glial fibrillary acidic protein (GFAP, green) at the site of injury, assessed at 1 day post-injury (dpi) and 3 dpi following different treatments under AMF (Scale bar = 50 μm). **c**, Quantitative analysis of the corrected total intensity of GFAP in various experimental groups at 1 dpi and 3 dpi (n = 6, means ± s.e.m.). **d**, Mean distance from the injury site observed in different treatment groups at 1 dpi and 3 dpi. For each fish, measurements were taken at five equidistant points from the top to the bottom of the injury site, and the average of these measurements was calculated (n = 6, means ± s.e.m.). **e**, Live imaging of the colocalization of GFAP in a transgenic line with NPC/NPCbot cells labeled with CellTracker Deep Red and the corresponding scatter plot with Pearson’s correlation coefficient (Scale bar = 40 μm). **f**, Quantitative evaluation of colocalization, employing Pearson’s correlation coefficient, across five replicates (n = 5, means ± s.e.m.). Statistical significance was calculated via one-way ANOVA with a Tukey post-hoc test (**Fig. 4c, d, f**).* p < 0.05, **** p< 0.0001 versus control.

Further supporting the role of NPCbots in engaging astrocytes in the neural regeneration process, we tracked how effectively our NPCbot treatment integrated into this spinal repair mechanism. Additional analyses involved live imaging in conjunction with a zebrafish transgenic line labeling GFAP to monitor astrocyte dynamics. We tracked the progression of regeneration with and without NPCbot treatment in the same fish, exploring the comparative impact of our intervention on the regeneration process (**Fig. 4e**). When comparing the NPCbot group to the control group treated with NPCs only, we observed a 2.6-fold increase in the colocalization of GFAP expression and NPC/NPCbot cells labeled with CellTracker Deep Red by day three post-injury (3 dpi). This colocalization, evident in the merged yellow fluorescence in the images, underscored an enhanced integration of differentiated NPCbots with host astrocytes. The Pearson’s correlation coefficient, a measure of colocalization, indicated a 2-fold increase at 3 dpi in the NPCbot-treated group compared to the controls (**Fig. 4f**). This finding confirms that NPCbots not only differentiate into astrocytic lineages expressing GFAP but also actively incorporate into the healing spinal tissue, thereby potentially facilitating more effective regenerative processes. Our findings unequivocally demonstrate that NPCbot treatment, particularly when combined with CFO-BTO and stimulated by AMF, significantly enhances neural connection and spinal cord regeneration in a zebrafish model. The observed improvements in neural and astrocytic marker expression, along with the physical contraction of the injury site, suggest that NPCbots can effectively integrate into and support the regenerative milieu.

### Behavioral Assessment and Recovery in NPCbot-Treated Zebrafish

In this segment of our study, we carried out extensive behavioral assessments to ascertain both structural and functional recovery in zebrafish following spinal cord injury and subsequent treatment with NPCbots. Utilizing a 96-well plate, we divided the wells to accommodate six zebrafish groups subjected to different treatment protocols, with each group allocated 16 wells. Observations spanned a 15-minute period, segmented into alternating light and dark cycles of five minutes each, to gauge zebrafish responses to environmental light shifts (visual motor response) – a vital parameter for assessing the efficacy of NPCbot interventions on zebrafish behavior. We measured swimming distance and speed – both average and peak – alongside movement patterns.

At 1 dpi, there was a notable disparity between the healthy (G1, G2) and injured groups (G3-G6). Unlike the injured groups, which exhibited significantly reduced movement or paralysis in more severe cases (**Fig. 5a-e**). This diminished mobility underscores the critical role of the spinal cord in relaying movement signals from the brain to the body. Moreover, injured fish demonstrated general impairments in swimming behavior under varying light conditions and did not show noticeable motion, as indicated by their trajectories in **Fig. 5b**. Heat map analysis further delineated these behavioral modifications, accentuating areas within the wells where zebrafish lingered most frequently (**Fig. 5c**). By day 3 (1 dpi), healthy zebrafish displayed increased activity compared to the other groups, with movement patterns that suggest broader mobility within their designated wells. Conversely, those suffering from spinal injuries displayed a stark contrast, predominantly remaining immobile and confined to limited areas, indicative of their compromised motor functions and altered exploratory behavior.

**Fig. 5.**
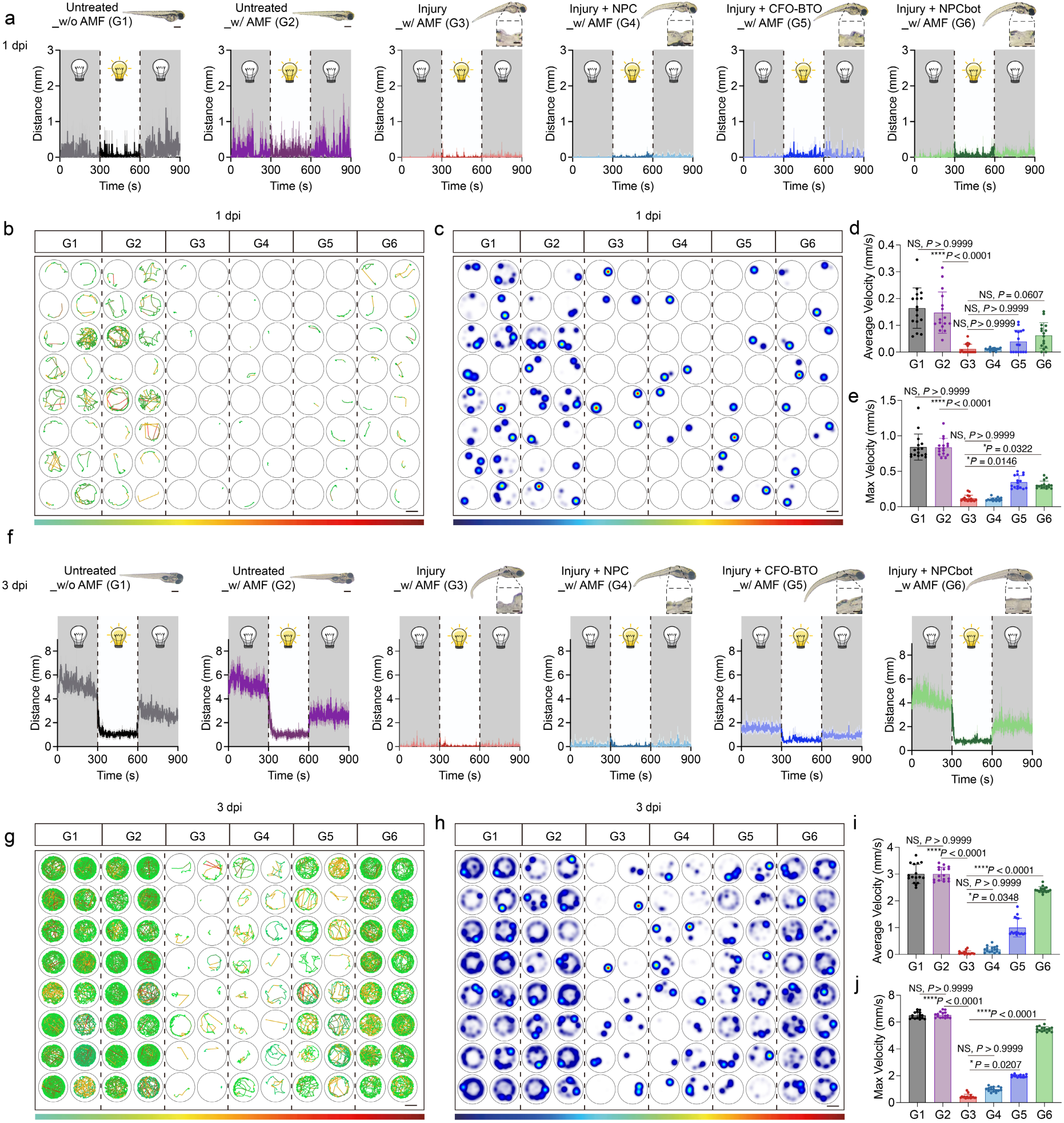
Behavioral Assessment and Recovery in NPCbot-Treated Zebrafish Larvae. **a**, Representative images of zebrafish larvae at 1-day post-injection (dpi) following different treatments. Swimming distance was assessed using a “dark-light-dark” protocol, each phase lasting 5 mins (Scale bar = 1 mm). **b**, Trajectory and **c**, heatmap illustrating the movement patterns of differently treated larvae (Scale bar = 4 mm). **d,** Quantitative analysis of the average and **e**, maximum swimming speeds of the treated larvae, respectively (n = 16, means ± s.e.m.). **f**, Representative images of zebrafish larvae at 1 dpi following different treatments. Swimming distance was assessed using a “dark-light-dark” protocol, each phase lasting 5 mins (Scale bar = 1 mm). **g,** Trajectory and **h**, heatmap illustrating the movement patterns of differently treated larvae (Scale bar = 4 mm). **i,** Quantitative analysis of the average and **j**, maximum swimming speeds of the treated larvae, respectively (n = 16, means ± s.e.m.). Statistical significance was calculated via Kruskal–Wallis non-parametric test with Dunnett’s post hoc analysis (**Fig. 5d, e, i, j**). * p < 0.05, **** p< 0.0001 versus control. For the trajectory analysis (**Fig. 5b,g**), the color red indicates an increased swimming speed, while green denotes a decreased speed. In the heatmap analysis (**Fig. 5c,h**), red signifies increased dwell time, and blue signifies decreased dwell time

Significant enhancement was evident by day 5 (3 dpi) in the NPCbot-treated group (G6), in comparison to those receiving only nanoparticle treatment (G5) or left untreated (G3). NPCbot-treated zebrafish not only demonstrated elevated overall swimming speeds and distances but also displayed movement difference in response to different light conditions, indicative of a resurgence in neural function (**Fig. 5f-j**). The behavioral data extracted from the Zebrabox, an automated observation chamber and behavioral tracking system for zebrafish, unequivocally showed marked improvements in motor function, not only facilitated structural repair but also restored functional behaviors. Further analysis of swimming patterns showed a tendency among the NPCbot-treated zebrafish to swim along the tank edges in circles, a behavior suggesting enhanced curiosity and improved spatial awareness (**Fig. 5h**). This behavior not only mirrors better motor function but also hints at a recovery in sensory and neural processing capabilities, facilitating more effective interaction with their surroundings. Additionally, it is noteworthy that NPC treatment alone (G4) did not significantly enhance the recovery of motor behaviors in injured zebrafish.

Together, these detailed behavioral assessments and quantitative analyses substantiate that NPCbot treatment significantly aids in the recuperation of motor functions and behavioral patterns in zebrafish post-spinal cord injury. The reinstatement of almost normal swimming behavior, and improved exploratory behaviors in NPCbot-treated zebrafish underscore the potential of this innovative treatment strategy in fostering comprehensive neural regeneration. These findings not only emphasize the efficacy of NPCbots in structural neural repair but also highlight their role in the functional restoration of the central nervous system.

### Western Blot Analysis and Mechanistic Insights into NPCbot-Induced Neural Regeneration

To elucidate the molecular mechanisms underlying NPCbot-induced neural regeneration, we employed Western blotting to quantitatively assess protein expression levels of key neuronal and astrocytic markers at the cellular level. βIII-Tubulin and MAP2, pivotal for the structural integrity and connectivity of neurons, were examined. βIII-Tubulin serves as an indicator of neuronal formation and growth, while MAP2 enhances the stability and linkage among neuronal branches. Our results demonstrated significant increases in these proteins of NPCbots group under an AMF, highlighting not only neuronal growth but also the facilitation of neuronal regeneration (**Fig. 6a-f**). These molecular changes are indicative of the potential for improved motor functions once translated to an *in vivo* model. Further, we analyzed the expression of GFAP to understand the response of astrocytes – supportive glial cells essential for neural repair. The expression levels of GFAP indicated that the NPCbot group optimally activates astrocytes to aid in neural repair. Additionally, α-Tubulin, known for its role in maintaining neuronal shape and intracellular transport, showed increased abundance in the NPCbot group under AMF (**Fig. 6g-l**). This suggests that our treatment supports the structural rebuilding and functional improvement of neurons, potentially contributing to enhanced locomotive abilities once applied *in vivo*.

**Fig. 6.**
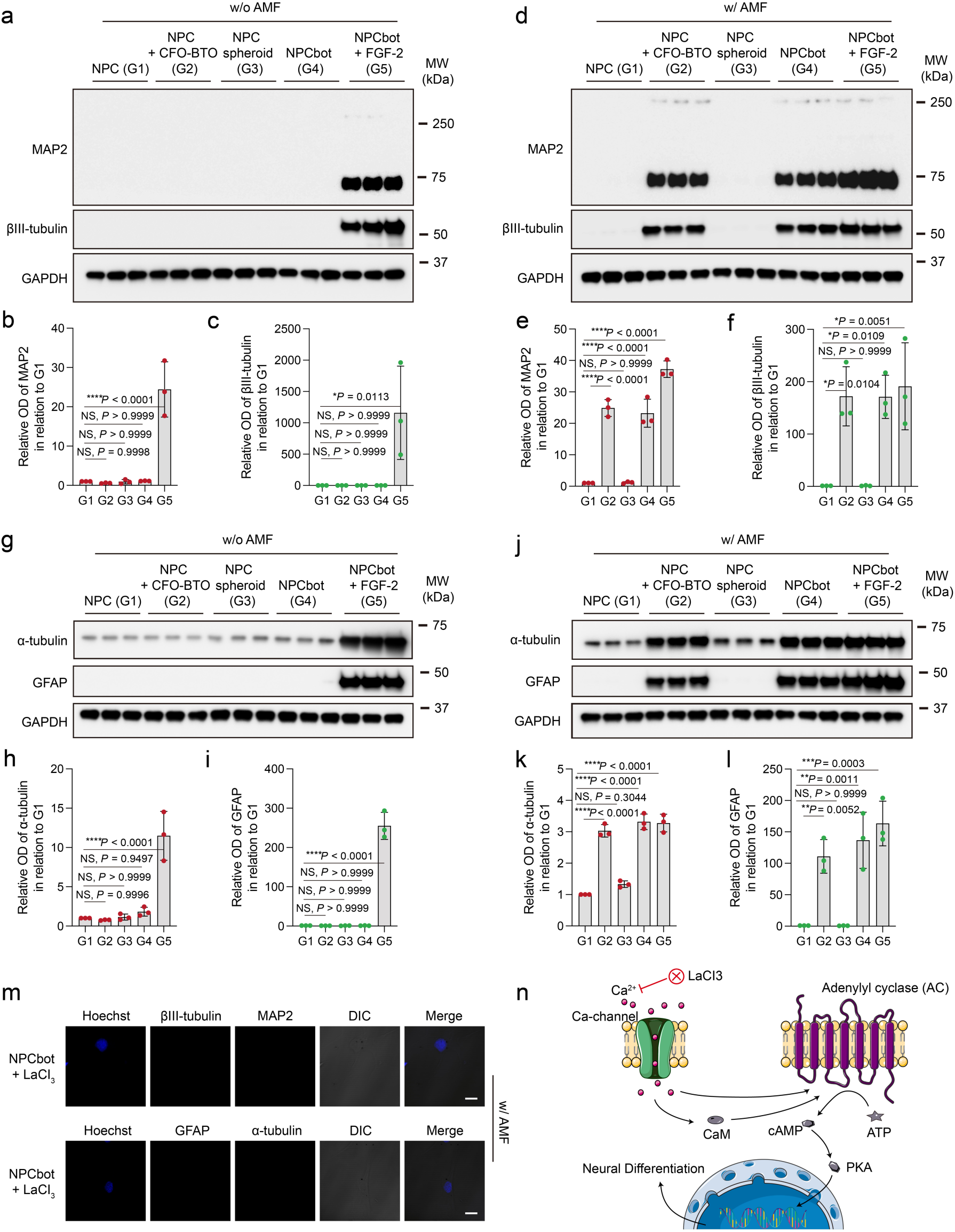
Western Blot Analysis and Mechanistic Insights into NPCbot-Induced Neural Regeneration. **a**, Western blot assays and quantitative analysis displaying the expression levels of neuron-specific markers. **b**, Relative quantification of MAP2 and **c,** βIII-tubulin expression following various treatments (n = 3, means ± s.e.m.). **d-f**, Western blot analysis of MAP2 and βIII-tubulin expression under AMF after different treatments (n = 3, means ± s.e.m.). **g-i**, Western blot assays detailing the expression levels of the neurite marker α-tubulin and the astrocytic marker GFAP following various treatments (n = 3, means ± s.e.m.). **j-l**, Quantification of α-tubulin and GFAP expression under AMF post-treatment (n = 3, means ± s.e.m.). **m**, Inhibitor experiments showcasing immunofluorescence analysis of NPCbots, indicating the expression of various differentiation markers stimulated by AMF in the presence of LaCl_3_ inhibitor (Scale bar = 8 μm). **n**, Schematic depiction of the intracellular pathways implicated in NPC differentiation influenced by NPCbots and external AMF stimulation. Statistical significance was calculated via one-way ANOVA with a Tukey post-hoc test (**Fig. 6b, c, e, f, h, i, k, l**).* p < 0.05, ** p < 0.01, ***p< 0.001, **** p< 0.0001 versus control.

To further elucidate the differentiation mechanisms of NPCs under NPCbot treatment, our research focused on the cAMP-dependent pathway, also known as the adenylyl cyclase (AC) pathway^35^. This pathway can be activated by various extracellular stimuli^15, 36^. Elevated intracellular Ca^2+^ concentrations, whether directly or indirectly, form a complex with calmodulin (CaM), leading to the activation of AC^37^. AC is a pivotal regulatory enzyme that catalyzes the conversion of ATP (Adenosine triphosphate) to cyclic AMP (cAMP). Injured fish demonstrated impaired responses to changes in environmental light shifts. The second messenger cAMP activates the protein kinase A (PKA)-dependent pathway for neural progenitor cell differentiation.^38^. Prior research indicates that electrical stimulation can activate intracellular Ca^2+^ channels, resulting in an increased influx of intracellular Ca^2+39, 40^. We hypothesized that both electrical and magnetoelectric stimulations, as delivered by the AMF in our NPCbot system, enhance calcium influx through these channels, thereby amplifying the cAMP-dependent differentiation pathway. To validate this hypothesis, we introduced LaCl_3_, a calcium channel blocker^41^, into our experimental set-up. Utilizing confocal microscopy for in-depth analysis, we observed a marked decrease in the expression of both neuronal and astrocytic markers upon the addition of LaCl_3_ (**Fig. 6m, n** and **Supplementary Fig. 6**). This finding underscores the crucial role of calcium channel activation in the AMF-dependent differentiation process facilitated by NPCbots. These insights not only provide a molecular basis for the observed enhancements in structural and functional cellular attributes but also substantiate the mechanistic efficacy of our NPCbot treatment.

### *In vitro* and *in vivo* NPCbot manipulation

To evaluate the translational potential of NPCbot for clinical applications, we explored their navigation capabilities both *in vitro* and *in vivo*. Utilizing a state-of-the-art magnetic manipulation system (MFG-100, MagnebotiX AG, Switzerland) capable of five-degrees-of-freedom (5-DOF) wireless micromanipulation (**Fig. 7a**), we assessed the motility of NPCbots within a phosphate-buffered saline (PBS) solution. The results, illustrated in **Fig. 7b,c** and further illustrated by videos in **Supplementary Video 1**, demonstrate that under a rotating magnetic field of 10 mT at a frequency of 1 Hz, NPCbots can execute a rolling motion to follow a predefined trajectory – spelling “ETH” in this instance – within a glass petri dish. The NPCbots displayed reliable magnetic responsiveness, navigating the set path with minimal deviation, which highlights their potential for precise path-following capabilities in fluidic environments.

**Fig. 7.**
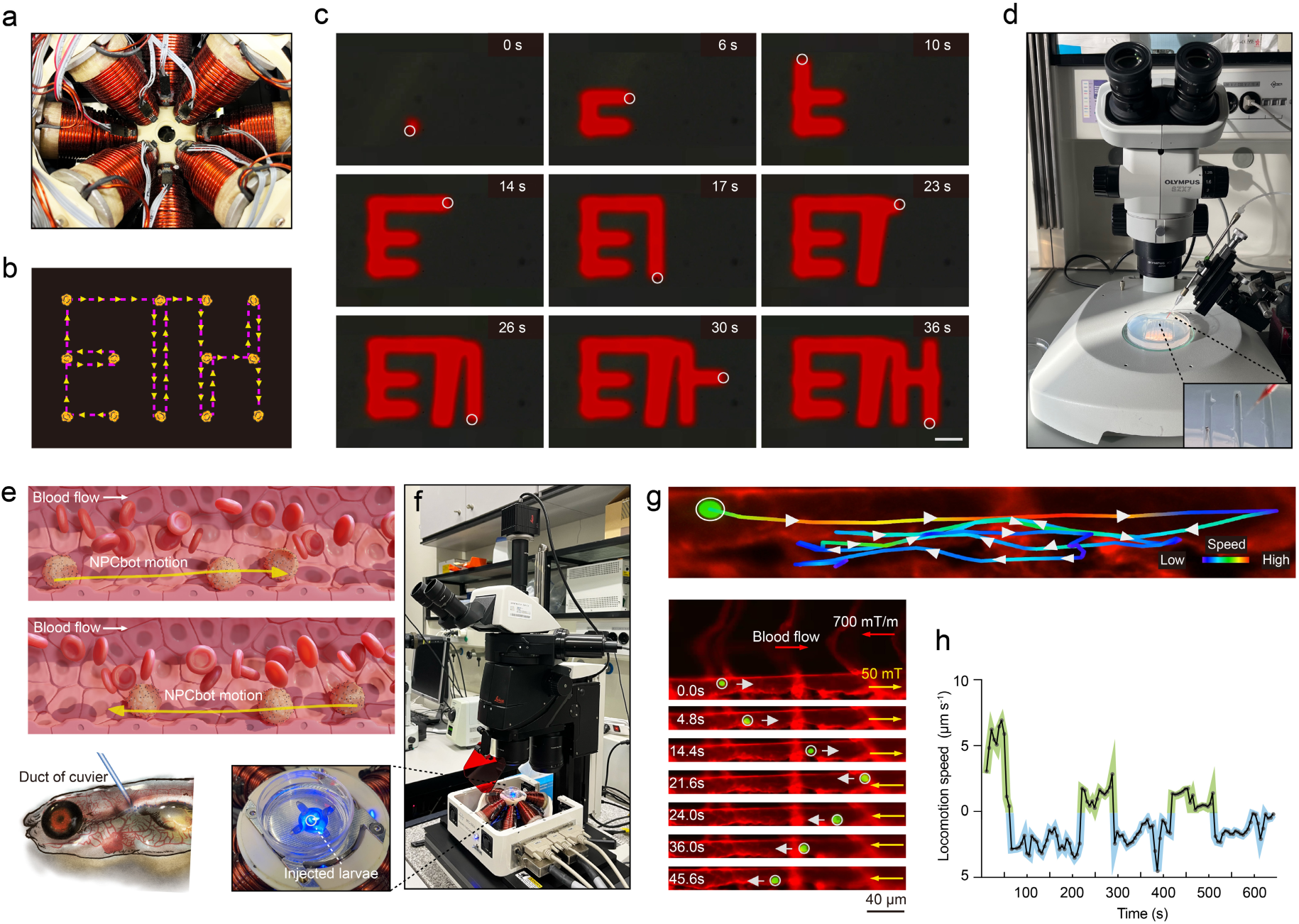
*In Vitro* and *In Vivo* Manipulation of NPCbots. **a,** Photograph of the magnetic manipulation apparatus, which enables five degrees of freedom (5-DOF) for wireless micromanipulation. **b,** Preset motion trajectory for the NPCbot. **c,** Microscope images illustrating the locomotion of magnetic rolling NPCbots under an external rotating magnetic field of 10 mT at 1 Hz in a phosphate-buffered saline (PBS) solution, with the NPCbots’ positions highlighted by white circles (Scale bar = 20 μm). **d,** Photographs of the injection setup. **e-f,** Schematic of the *in vivo* manipulation setup and navigation scenario following injection into zebrafish larvae, showing how the NPCbot can navigate both with and against the blood flow. **g,** Trajectory analysis of NPCbot movement within the dorsal aorta (DA) of kdrl:mCherry zebrafish larvae, highlighted by white arrows indicating the direction of travel. The NPCbots, labeled with live dye in green, are depicted navigating both with and against the blood flow. This is achieved by applying compensatory magnetic field gradients of 50 mT and 700 mT/m to counteract the hemodynamic forces. The stereomicroscope images (below) capture the real-time positioning of the NPCbots within the fluorescently highlighted vascular architecture (Scale bar = 40 μm). **h)** Analysis of NPCbot locomotion speed within the DA of zebrafish larvae, as measured using Imaris software. The green line represents the NPCbot movement in the same direction as the blood flow (co-directional navigation), and the blue line represents movement opposite to the blood flow (counter-directional navigation).

Transitioning to *in vivo* experiments, NPCbots were pre-labeled using a Live dye (L3224, LIVE/DEAD™ Viability/Cytotoxicity Kit, Thermo Fisher Scientific) and introduced into the duct of Cuvier in zebrafish embryos at 2 dpf by injection of 2 to 3 nL of NPCbot solution using a FemtoJet injection system (Eppendorf) as depicted in **Fig. 7d**. A rotating magnetic field of 50 mT at 3 Hz, coupled with a compensatory magnetic field gradient of 700 mT/m, was employed to counteract the influence of blood flow within the dorsal aorta (DA), a major vessel in zebrafish embryos characterized by high flow rates. This setup allowed for precise manipulation of NPCbots within the vascular system. The *in vivo* locomotion of the NPCbots was meticulously documented utilizing a Leica M205 FCA stereomicroscope, as depicted in **Fig. 7e, f**. To further scrutinize the movement dynamics of the NPCbots *in vivo*, we employed the kdrl:mCherry transgenic line to visualize blood vessels, while the NPCbots were distinctly labeled with Live dye (FITC channel). Our comprehensive analysis, illustrated in **Fig. 7g**, demonstrates that the NPCbots could be manipulated within the designated area. The NPCbots performed controlled back-and-forth movements (**Supplementary Video 2**), spanning a distance of 112 µm. Despite the dynamic environment posed by the blood flow within the DA, the NPCbots traveled a maximum distance of 156 μm from their initial position, indicating effective navigational control.

Furthermore, the locomotion of NPCbots within the DA was systematically analyzed over a 600-second interval, as depicted in **Fig. 7h**. When moving with the blood flow (green curve), NPCbots occasionally reached peak speeds of nearly 8 µm/s, likely benefiting from the additive velocity of the blood flow. However, these peak speeds were not consistently maintained. Conversely, the movement against the blood flow (blue curve) showcased generally lower speeds but greater stability. This pattern of movement indicates a consistent resistance from the opposing blood flow, which limits peak speeds but results in a steadier navigation pattern. The navigation analysis underscores the capabilities of NPCbots for targeted navigation and actuation within both *in vitro* and *in vivo* environments. The precise control achieved over NPCbot movement, especially within complex biological systems like the zebrafish vascular network, highlights the significant potential of magnetic microrobots in therapeutic applications, including targeted drug delivery and regenerative medicine.

## CONCLUSIONS

The development and application of NPCbots described in this study represent an advance in the field of regenerative medicine, especially in the treatment of spinal cord injuries. By integrating microfluidics technology with magnetoelectric nanoparticles and employing real-time magnetic navigation, this approach introduces a novel method for precision therapeutic delivery and cellular manipulation. This innovative approach may influence transform treatment paradigms not only for spinal cord injuries but also for a broader spectrum of neurodegenerative diseases.

The positive results observed in zebrafish models indicate that magnetic navigation combined with *in vivo* stimulation has significant potential in regenerative medicine. As we continue to refine these technologies, NPCbots could play a valuable role in the treatment of various neurological disorders and injuries, providing targeted, efficient, and minimally invasive interventions. Moreover, the conceptual and technological frameworks developed in this study may inspire similar approaches in other medical fields. The principles of targeted delivery and controlled therapeutic action demonstrated by NPCbots are likely to be beneficial in specialties such as oncology and cardiology, where precise localization and controlled drug release are crucial.

In future applications, the NPCbot platform is set to advance cell therapy by significantly enhancing cell survival, precision targeting, and *in vivo* stimulation capabilities. Nevertheless, this path is not without its challenges and opportunities for further exploration. Future research should focus on enhancing the scalability of NPCbot production to facilitate broader clinical applications. This entails optimizing the LoC designs to increase throughput and efficiency without compromising the functionality and integrity of the bots. Additionally, investigating various cell types and nanoparticle formulations could lead to better outcomes or adaptable strategies for treating a range of neurological conditions.

Importantly, further studies are needed to evaluate the long-term viability and safety of NPCbots in larger animal models. These studies are vital for gathering data on the immunological impacts, potential toxicity, and persistent effects of nanoparticle integration – key factors for advancing from zebrafish models to human clinical trials. Extending *in vivo* testing to mammalian models will be essential for understanding the complex interactions within a more human-like neurological setting, potentially unveiling new challenges or necessary adjustments for human application. Concurrent technological advancements in imaging and real-time tracking are imperative to support the detailed monitoring and precise control of NPCbots within the body. Moreover, enhancing NPCbots with real-time feedback mechanisms could lead to adaptive treatment strategies, allowing the bots to modify their therapeutic actions dynamically in response to the evolving microenvironment at the target site.

## Supporting Information

Supporting Information is available online or from the author.

## Acknowledgements

The authors thank David Sargent for his help with the magnetic manipulation test. This work was financially supported by the Swiss National Science Foundation (No. 206033) and National Natural Science Foundations of China (Grant 82161138029) for a Sino-Swiss Science and Technology Cooperation project, National Natural Science Foundation of China (No. 82073777). The authors would also like to acknowledge the financial support from the Swiss National Science Foundation under project number No. 198643, European Union’s Horizon Europe Research and Innovation Programme under EVA project (GA No: 101047081) and META-BRAIN project (GA No: 101130650). The Swiss State Secretariat for Education, Research and Innovation (SERI) is also acknowledged. The authors would also like to thank the Scientific Center for Optical and Electron Microscopy (ScopeM), and the FIRST laboratory at ETH for their technical support.

## Data Availability Statement

All data associated with this study are present in the main text or the Supplementary Materials. Additional data is available from authors upon request.

