## Supporting Information for "Magnetoelectric Microrobots for Spinal Cord Injury Regeneration"

### EXPERIMENTAL SECTION

**Synthesis of CFO-BTO core-shell magnetoelectric nanoparticles.** Cobalt ferrite-barium titanate (CFO-BTO) core–shell nanoparticles were synthesized through a sequential application of co-precipitation, hydrothermal, and sol-gel methods, as detailed in prior studies[1]. Initially, 2 g of cetyltrimethylammonium bromide (CTAB) was dissolved in 30 mL of deionized (DI) water. Following this, 1 g of ferric chloride hexahydrate ($FeCl_3 \cdot 6H_2O$) and 0.24 g of anhydrous cobalt(II) chloride ($CoCl_2$) were separately dissolved in the CTAB solution. A 6 M sodium hydroxide (NaOH) solution was then gradually added to initiate the precipitation of CFO nanoparticles. For the transformation of the CFO nanoparticles to a single-crystalline structure, the resultant mixture was sealed in an autoclave and subjected to a hydrothermal treatment at elevated temperatures. Subsequently, the formation of the BTO shell over the CFO core was achieved using a sol-gel process. The BTO precursor was prepared by combining 30 mL of a DI water solution containing 0.029 g of barium carbonate ($BaCO_3$) and 0.1 g of citric acid with 30 mL of an ethanolic solution, which included 1 g of citric acid and 0.048 mL of titanium isopropoxide ($Ti(OCH(CH_3)_2)_4$). This mixture was continuously stirred at 90 °C for 1 hour to ensure complete sol formation. CFO nanoparticles, previously dispersed via sonication for 2 hours, were added to the sol to form a gel. This gel was dried at 80 °C overnight. The dried gel was subsequently annealed at 600 °C to yield CFO-BTO nanoparticles.

**Characterization of CFO-BTO core-shell magnetoelectric nanoparticles.** The morphology of the nanoparticles was examined using scanning transmission electron microscopy (STEM, FEI Talos F200X). The elemental distribution within the nanoparticles was assessed by high-angle annular dark field (HAADF) STEM on a FEI Talos F200X system with energy-dispersive X-ray (EDX) spectroscopy for elemental mapping. X-ray diffraction analyses were performed using an Empyrean diffractometer, which utilizes a copper X-ray source ($\lambda$ = 1.5406 Å) and is coupled with a PIXcel detector (Malvern Panalytical). Local piezoresponse loops were evaluated using piezoresponse force microscopy (PFM, NT-MDT Ntegra Prima). A conductive AFM tip (FMG01/Pt) in contact mode applied an alternating voltage of 0.5 V to the CFO-BTO NP, inducing piezoelectric surface oscillations. These oscillations were detected via cantilever deflection. Local piezoresponse hysteresis loops were generated at random NP locations by modulating the applied DC bias and simultaneously recording the phase and amplitude responses. The excitation voltage was a stepwise increasing pulsed DC voltage, augmented with a small AC voltage to mitigate electrostatic interference, with the AC response signal captured exclusively during the off-phase of the pulse sequence. To investigate changes in the piezoelectric response under magnetic influence, an in-plane magnetic field of 50 mT was applied to the sample.

**Human iPSC-derived neural progenitor cell (NPC) culture.** In the protocol for cultivating human induced pluripotent stem cell (iPSC)-derived neural progenitor cells (NPCs), the utilization of Matrigel-coated culture vessels was crucial. The process commenced with the careful thawing of Matrigel at reduced temperatures, followed by a dilution at a ratio of 1:50 with chilled Dulbecco's Modified Eagle Medium (DMEM). This mixture was subsequently applied to culture flasks, which were then incubated overnight within a temperature range of 4 °C. Prior to cell seeding, the coating solution was aspirated and replaced with pre-warmed ENStem-A Neural Expansion Medium (Catalogue No. SCM004a, Merck). Cryopreserved human iPSC-derived NPCs (Catalogue No. SCC035, Merck) were swiftly thawed at 37 °C in a water bath, then transferred into a sterile 15 mL conical tube. Here, they were gently mixed with pre-warmed medium to mitigate osmotic shock. The cell suspension was subsequently centrifuged, the supernatant decanted, and the cell pellet resuspended. Cell viability was assessed through counting before seeding onto the prepared Matrigel-coated culture vessels. Cultures were maintained at 37 °C in a humidified atmosphere containing 5% CO2, with medium exchanges every 2 days. Upon attaining confluence, NPCs were dissociated using Accutase (Catalogue No. SCR005, Merck), quantified, and replated into similarly prepared T75 flasks. This process promoted optimal conditions for the proliferation and maintenance of neural progenitor cells.

**Fabrication and characterization of NPCbot.** Microfluidic chips were fabricated using replica molding technique. Initially, master molds were created on silicon wafers employing photolithography techniques and wet etching, utilizing a negative photoresist (SU-8 3010, Micro Resist Technology GmbH). Following this, polydimethylsiloxane (PDMS) slabs were prepared over the silicon master molds through the replica molding process. For this step, 10:1 PDMS:curing agent (The Dow Chemical Company) mixture was prepared and degassed in the desiccator for 30 min. Then the mixture was poured over the silicon master molds and cured in the oven at 60 °C for 45 min. After curing process, PDMS slabs were precisely cut and punched to establish inlet and outlet holes utilizing a 1.5 mm biopsy puncher. Subsequently, these PDMS chips were irreversibly bonded to a PDMS-coated glass slide via oxygen plasma activation to complete the fabrication process.

During the microrobot fabrication phase, cells were isolated within the traps located in the trapping chamber of the lab-on-a-chip (LoC) device. This isolation was achieved via flushing the growth medium solution including NPCs (at a concentration of $5\times10^5$ cell/mL) at a flow rate of 1 μL/min for 20 min, which was then followed by the flow of magnetoelectric nanoparticle solution in growth medium (33 mg/mL) for *in situ* NPCbot incubation. After the 30 min incubation period, the outlet of the LoC device was reconfigured as an inlet for the reverse flow (growth medium at 15 μl/min), facilitating the release of the incubated NPCbots from the traps, while the three inlets were utilized as outlets for collecting the NPCbots. This bidirectional design of the LoC device enhances the efficiency of the trapping and release mechanisms, thereby supporting the mass production and expedited *in situ* fabrication of biohybrid microrobots, *i.e.* NPCbots. All experimental procedures were meticulously documented using an Olympus IX73 microscope.

To assess the viability of the NPCbots, we employed live/dead staining. An NPC cell stock solution, at a concentration of 1000 cells/μL, was incubated with CFO-BTO nanoparticles at varying concentrations (0, 500, 1000, 1500, 2000 ng/μL) for a duration of 2 hours to obtain different concentrations of NPCbots. The viability was subsequently evaluated using the Live/Dead Cell Imaging Kit (488/570) from Invitrogen, USA. This kit utilizes a combination of Live Green and Dead Red components, which were mixed and incubated for 15 minutes at room temperature. Fluorescence images were captured using a confocal microscope (LSM 880; Carl Zeiss, Germany) and analyzed with ImageJ software. This procedure facilitated the determination of an optimized CFO-BTO dosage, which was then used to ascertain the ideal incubation time needed to maintain the nanoparticles on the cell surface, thereby minimizing cellular uptake.

To further investigate the cellular uptake of CFO-BTO nanoparticles, we labeled them with Alexa Fluor 594. This labeling involved a two-step surface modification process: First, CFO-BTO nanoparticles were coated with (3-Aminopropyl)triethoxysilane (APTES) to introduce amino groups on their surface. This was performed by dispersing the nanoparticles in a 5% (v/v) APTES ethanol solution and allowing the reaction to proceed with continuous stirring at room temperature for 2 hours. Post-APTES coating, the nanoparticles were centrifuged at 10,000 g for 10 minutes, washed three times with ethanol, and redispersed in phosphate-buffered saline (PBS, pH 7.4). Fourier Transform Infrared Spectroscopy (FTIR) was used for characterization. For the conjugation of Alexa Fluor 594, the APTES-coated nanoparticles were incubated with Alexa Fluor™ 594 NHS Ester (Succinimidyl Ester) at a concentration of 0.1 mg/mL in PBS. This step, conducted at room temperature for 1 hour under gentle agitation, allowed the NHS ester to react with the amine groups to form stable amide bonds. The nanoparticles were subsequently centrifuged, washed three times with PBS to remove excess dye, and resuspended in PBS. They were stored at 4 °C and protected from light until further use. To quantitatively assess the cellular uptake, dye-labeled CFO-BTO nanoparticles were incubated with cells for 0.25, 0.5, 1, and 2 hours. Following incubation, cells were washed eight times with cold PBS and analyzed using flow cytometry (LSRFortessa, BD Biosciences). A Hitachi SU5000 field-emission scanning electron microscope (SEM) was employed to examine the morphology of the NPCbot. Additionally, the distribution of CFO-BTO on the NPC surface was analyzed using the Zeiss LSM 880 with Z-stack analysis.

**Stimulation of NPCbot by an alternating magnetic field.** For the assessment of the differentiation of NPCbot, the samples were subjected to an alternating magnetic field (AMF) stimulation protocol. AMFs were applied using the setup consists of a solenoid coil with inside diameter of 5 cm in which the samples were placed. The solenoid was driven in a resonant tank configuration and generated a magnetic field of amplitude 20 mT and frequency 1.18 kHz at its center. Air-cooling was used to stabilize the temperature of the solenoid. Each sample received AMF stimulation for two hours, twice daily over a two-day period. Following stimulation, the cells were fixed using 4% paraformaldehyde (PFA) for 10 minutes at room temperature. Permeabilization was facilitated by treating with 0.5% Triton X-100 for 30 minutes at room temperature. To minimize non-specific binding, the samples were blocked with 2% bovine serum albumin (BSA) for 30 minutes. For the detection of neuronal markers, the cells were incubated overnight at 4 °C with primary antibodies directed against βIII-tubulin (Sigma-Aldrich, T2200), microtubule-associated protein 2 (MAP2) (Proteintech, 67015-1), α-tubulin (Merck, T6793), and Glial Fibrillary Acidic Protein (GFAP) (Agilent, Z0334). This was followed by incubation with secondary antibodies: FITC-conjugated goat anti-rabbit IgG (Sigma-Aldrich, AP132F) and TRITC-conjugated goat anti-mouse IgG (Sigma-Aldrich, AP503R), each for 1 hour. After secondary antibody application, the samples underwent several washes and were subsequently stained with nuclear markers. Prior to NPC cell bot fabrication, immunofluorescence (IF) was utilized to verify the specificity of NPC cell markers, including Nestin (Sigma, ABD69) and SOX-2 (Sigma, MAB4343), using the aforementioned secondary antibodies. For the inhibitor experiments, the culture medium was supplemented with lanthanum chloride ($LaCl_3$) at a concentration of 50 μM. It is important to note that inhibitor were introduced to the culture medium prior to the commencement of the stimulation experiments. The stained samples were visualized using a Zeiss LSM 880 Airyscan confocal laser scanning microscope. Image quantification was performed using ImageJ software.

**Western blot analysis.** Protein extraction from cells, following exposure to an alternating current (AC) magnetic field, was conducted using Thermo Scientific™ RIPA Lysis and Extraction Buffer (Catalog number: 89900), which was supplemented with PMSF Protease Inhibitor (Catalog number: 36978) to ensure comprehensive protein recovery and inhibition of protease activity. Protein concentrations were determined utilizing the Thermo Scientific™ Pierce™ BCA Protein Assay Kit (Catalog number: 23227). Proteins were subsequently resolved by electrophoresis on SDS-PAGE gels, prepared using Tris-Glycine-SDS Buffer (Sigma, T7777) at the appropriate working concentration. Post-electrophoresis, proteins were transferred onto polyvinylidene fluoride (PVDF) membranes using Thermo Scientific™ Pierce™ 10X Western Blot Transfer Buffer, Methanol-free (Catalog number: 35040). To prevent nonspecific binding, membranes were blocked with Thermo Scientific™ Pierce™ Clear Milk Blocking Buffer (10X), diluted to a 1X concentration (Catalog number: 37587), and incubated for 1 hour at room temperature. Membranes were then incubated overnight at 4 °C with primary antibodies: Anti-beta III Tubulin (abcam, ab18207), Anti-alpha Tubulin (abcam, ab52866), Anti-GFAP (abcam, ab68428), and Anti-MAP2 (abcam, ab32454), all diluted following the manufacturer's recommendations. After primary antibody incubation, membranes were washed and incubated with HRP-conjugated Affinipure Goat Anti-Rabbit IgG(H+L) (Proteintech, SA00001-2) for 1 hour at room temperature. Detection of protein bands was achieved using Clarity™ Western ECL Substrate (Biorad, 200 ml #1705060) and visualized on an Amersham Imager 600 system, which was equipped with ImageQuant TL software for digital imaging and quantitative analysis.

**Zebrafish care.** Zebrafish (Danio rerio) were housed under controlled conditions, adhering to a 14-hour light/10-hour dark photoperiod, with illumination commencing at 8 am and concluding at 10 pm. Aquatic environments were consistently maintained within a temperature range of 26-28 °C[2]. Specimens from both the WIK wildtype strain and the GFAP transgenic line were employed in this investigation. Embryonic zebrafish were cultivated in E3 medium, which was composed of 5 mM NaCl, 0.17 mM KCl, 0.33 mM $CaCl_2$, and 0.33 mM $MgSO_4$. To prevent fungal contamination, the medium was supplemented with 0.01% methylene blue. Embryos were collected immediately post-oviposition and subsequently transferred to an incubation facility. This ensured that all larvae were reared under identical conditions with a stable ambient temperature set at 28 °C. All procedures involving live zebrafish were performed with approval from the Veterinary Authorities of Kanton Zurich, Switzerland, ensuring compliance with ethical standards for animal care and use.

**Manipulation of NPCbot via Magnetic Actuation:** A magnetic manipulation apparatus, facilitating 5-DOF (degrees of freedom) wireless micromanipulation, was utilized for evaluating NPCbot swimming capabilities (MFG-100, MagnebotiX AG, Switzerland), was employed to control the NPCbot's trajectory. These assessments were conducted in a phosphate-buffered saline (PBS) solution using a rotating magnetic field to propel the NPCbots. Movements were documented using an Olympus IX81 fluorescence microscope.

For *in vivo* manipulation within zebrafish larvae, NPCbots were pre-labeled using Live dye (L3224, LIVE/DEAD™ Viability/Cytotoxicity Kit, Thermo Fisher Scientific) as per the manufacturer's instructions. The NPCs were then introduced into the duct of Cuvier in zebrafish embryos (2 days post-fertilization, kdrl:mCherry line) embedded in low-melting-point agarose, using a FemtoJet injection system (Eppendorf). Each injection delivered a volume of 2 to 3 nl of NPCbot (200-300 NPCbot). Consistency in the amount of injected NPCbots was ensured by employing the same injection capillary for all larvae, with the process being monitored under a Leica M205 FCA stereomicroscope. Post-injection, the zebrafish embryos were carefully extracted from the original embedding medium and re-embedded in fresh low-melting agarose.

Magnetic control during *in vivo* experiments was achieved by applying a rotating magnetic field supplemented by a compensatory magnetic field gradient to counteract blood flow. The *in vivo* movements of NPCbots were recorded using a Leica M205 FCA stereomicroscope. Positional data were abstracted and analyzed using Imaris software (v9.9, Bitplane, Zurich, Switzerland), facilitating the calculation of swimming speeds, temporal data, and trajectory plotting of the NPCbots.

**Evaluation of NPCbot *In Vivo* Therapeutic Performance on Spinal Cord Injury in a Zebrafish larvae Model.** Following the assessment of NPCbot's mobility, its therapeutic efficacy was evaluated using a zebrafish spinal cord injury model. The NPCbots were injected at the site of injury and subjected to an alternating magnetic field. To establish the spinal cord injury model in zebrafish larvae, we anesthetized the larvae (2 days post-fertilization, dpf) in E3 medium supplemented with 0.02% Tricaine (ethyl 3-aminobenzoate methanesulfonate, MS-222). The anesthetized larvae were then transferred to a surgery plate. Excess E3 medium was carefully removed using an aspiration pipette. Positioned laterally on the plate, the larvae were approached dorsally with a microscalpel (30 G, 3 1/2-inch hypodermic needle, BD Biosciences, Cat#305106) modified by removing the barrel flange and attaching a cell scraper handle[3]. The incision was made across the dorsal trunk from the notochord to the dorsal edge, with the needle's bevel oriented sideways. Lesions were standardized to approximately one myotome in diameter (around 100 μm) while ensuring the notochord remained intact. These lesions were carried out under an optical microscope to confirm the complete excision of spinal cord tissue at the lesion site, maintaining consistency in the lesion location across all specimens. After lesioning, the larvae were gently transferred into a recovery dish using a pipette filled with E3 medium and subsequently placed in an incubator maintained at 28.5°C for recovery. The NPCbots were injected into the injury site, using 200-300 ng CFO-BTO and 200-300 NPCs as a control, followed by the application of an alternating magnetic field. Magnetic treatments were administered over a 2-hour period, with additional treatments provided on days 3 and 4 post-fertilization. Post-injection, the larvae were incubated at 34 °C.

The therapeutic efficacy of NPCbots at the molecular level was examined through whole-mount embryo immunohistochemistry. Larvae were pigment bleached, fixed in 4% formaldehyde in phosphate-buffered saline (PBS), and subsequently washed in PBS with 0.5% Tween and 0.5% Triton X-100 (PBST). Proteinase K treatment was conducted at a concentration of 10 μg/mL in PBST for 30 minutes at 37 °C, followed by a series of PBST washes and a post-fixation step in formaldehyde for 30 minutes at room temperature. Embryos were then treated with cold acetone at -20°C for 20 minutes, washed in PBST, and incubated in freshly prepared blocking solution (5% normal goat serum, 1% BSA, 1% DMSO in PBST) at room temperature for 2 hours. Primary antibodies specific for zebrafish GFAP (Agilent Cat#Z0334) and α-tubulin (Sigma-Aldrich Cat#T6793) were diluted 1:300 in the blocking solution and the embryos incubated overnight at 4 °C. Following this, embryos were washed in PBST and incubated overnight at 4 °C in the dark with secondary antibodies, Alexa Fluor 488 anti-rabbit (Thermo Fisher Scientific, Cat#A32731) and Alexa Fluor 555 anti-mouse (Thermo Fisher Scientific, Cat#A11008), both at a dilution of 1:300. After final washes in PBST, the embryos were embedded in 1% low-melting point agarose, positioned in confocal culture dishes (Thermo Fisher Scientific), and imaged using a Zeiss LSM 880 confocal laser scanning microscope. Image processing was performed using Fiji software.

To investigate whether NPCbots contribute to natural neural regeneration, live imaging techniques were employed. Specifically, the GFAP-transgenic zebrafish line expressing GFP was utilized. Following spinal cord injury (SCI) induction at 2 days post-fertilization (dpf), NPCbots (200-300 bots per specimen) were administered, with native neural progenitor cells (NPCs) serving as a control. Subsequent treatment involved the application of a uniform alternating current (AC) magnetic field for 2 hours per session on 2dpf, 3dpf, and 4dpf. Post-treatment, the zebrafish larvae were incubated at 34 °C. Imaging was performed using the LSM 880 confocal microscope, and the colocalization analysis was conducted using ImageJ software.

To further investigate the behavioral responses of zebrafish during our therapeutic intervention, we conducted assessments on six distinct groups that received treatment. The behavioral monitoring was executed using a Zebrabox (ViewPoint Life Sciences, Lyon, France). Each zebrafish larva was individually placed in a compartment within a 96-well plate, specifically designed to provide 16 separate Zebrabox environments. Observations were made over a 15-minute interval during which the larvae were subjected to alternating light and dark phases, each lasting 5 minutes. The movement of each larva was quantitatively recorded every second. All behavioral assessments were consistently performed within the same time period, specifically between 2 PM and 6 PM, at 1 and 3 days post-injury (dpi). The video footage captured during these sessions was subsequently analyzed with EthoVision XT11 software (Noldus Information Technology), which facilitated the generation of heat maps that depicted the positional distribution of the zebrafish over time.

**Statistical analysis.** All results were indicated as mean ± standard error of the mean (s.e.m.). One-way ANOVA (Tukey post-hoc tests) were applied for multiple comparisons and the Kruskal–Wallis non-parametric test with Dunnett's post hoc analysis was used to analysis the non-normally distributed data. All statistical analyses were conducted using the Prism software package (PRISM 9.5.0; GraphPad Software, 2022). $P < 0.05$ indicated statistical significance.

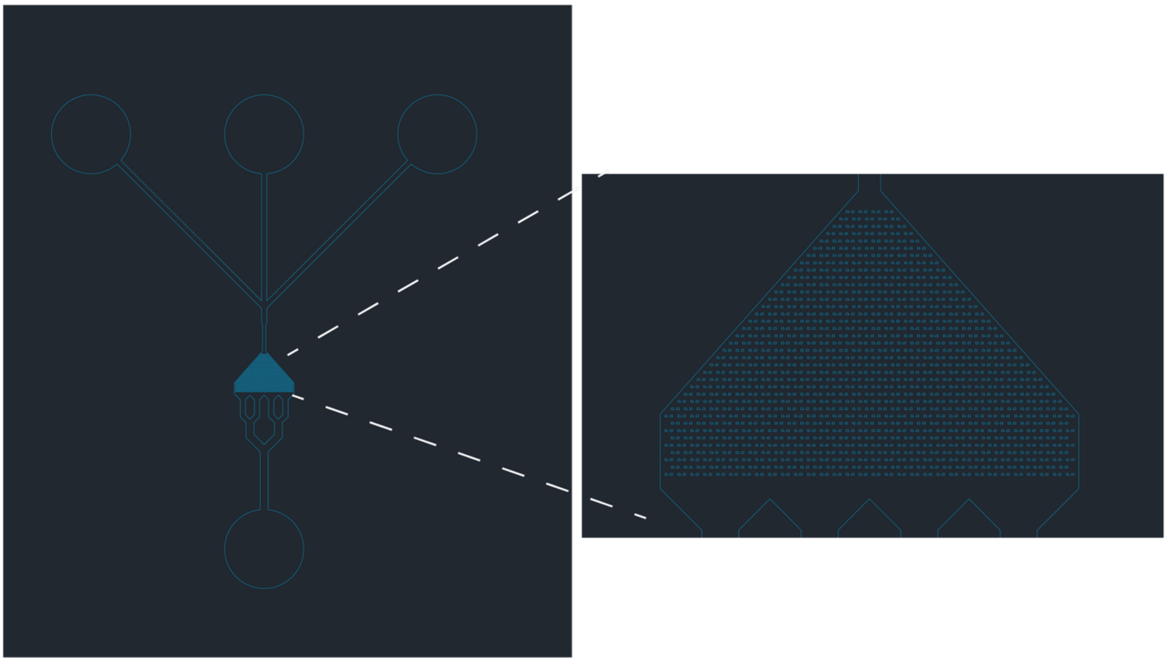

**Supplementary Fig. 1. Advanced Microfluidic Device for NPCbot fabrication**. This figure presents the schematic layout of a lab-on-a-chip system designed for precise cell manipulation and analysis. The inset provides a magnified view of the individual cell traps.

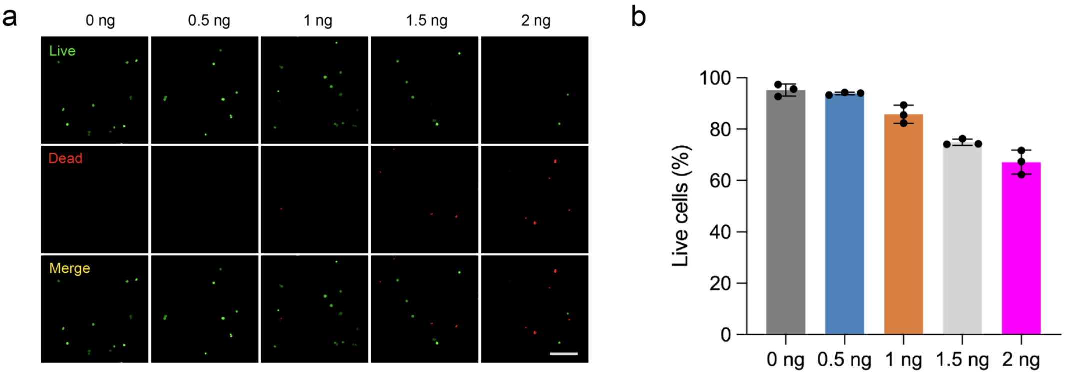


**Supplementary Fig. 2. Cell viability assessment of the fabricated-NPCbot across varying doses of CFO-BTO. a**, Live/dead assay and (**b**) MTT assay to optimize the CFO-BTO nanoparticle dosage required for NPCbot fabrication, indicating cell viability (green: live; red: dead; scale bar = 50 μm). As shown in the figure, less than 1 ng CFO-BTO per cell posses good biocompatatibility (n = 3, means ± s.e.m.).

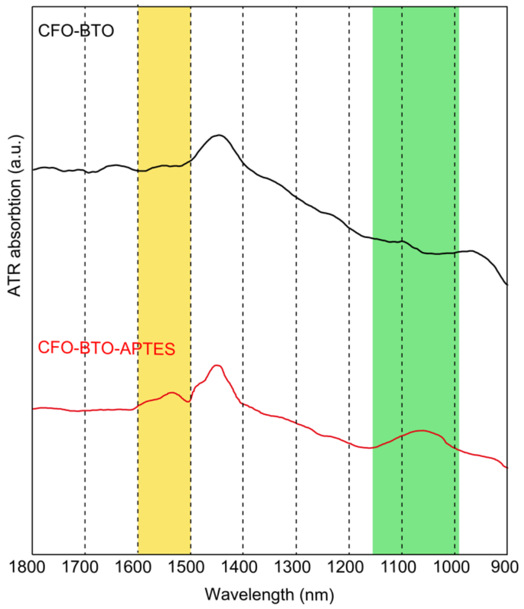


**Supplementary Fig. 3. FT-IR Spectra Analysis of Surface-Modified CFO-BTO.** This figure presents the FT-IR spectra for both non-coated and (3-aminopropyl)triethoxysilane (APTES)-coated CFO-BTO. The yellow highlighted region indicates the presence of amine groups, suggesting successful surface modification of CFO-BTO nanoparticles through APTES conjugation. Additionally, the green highlighted region shows the Si-O-Si signal, characteristic of APTES, confirming its conjugation to the CFO-BTO nanoparticles.

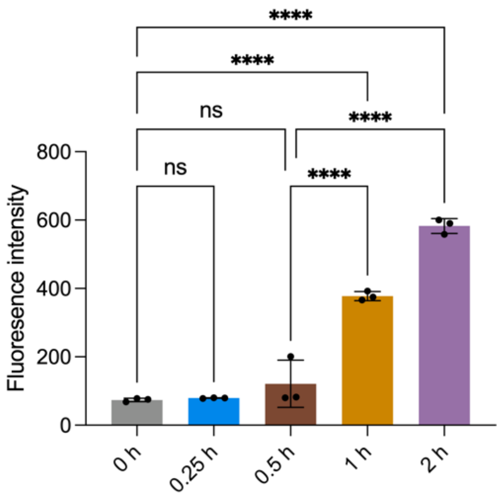


**Supplementary Fig. 4. Flow cytometry assays measuring the uptake of Alexa Fluor 594 by NPC following incubation periods of varying lengths.** n = 3, means ± s.e.m. Statistical significance was calculated via one-way ANOVA with a Tukey post-hoc test. **** $p < 0.0001$ versus control.

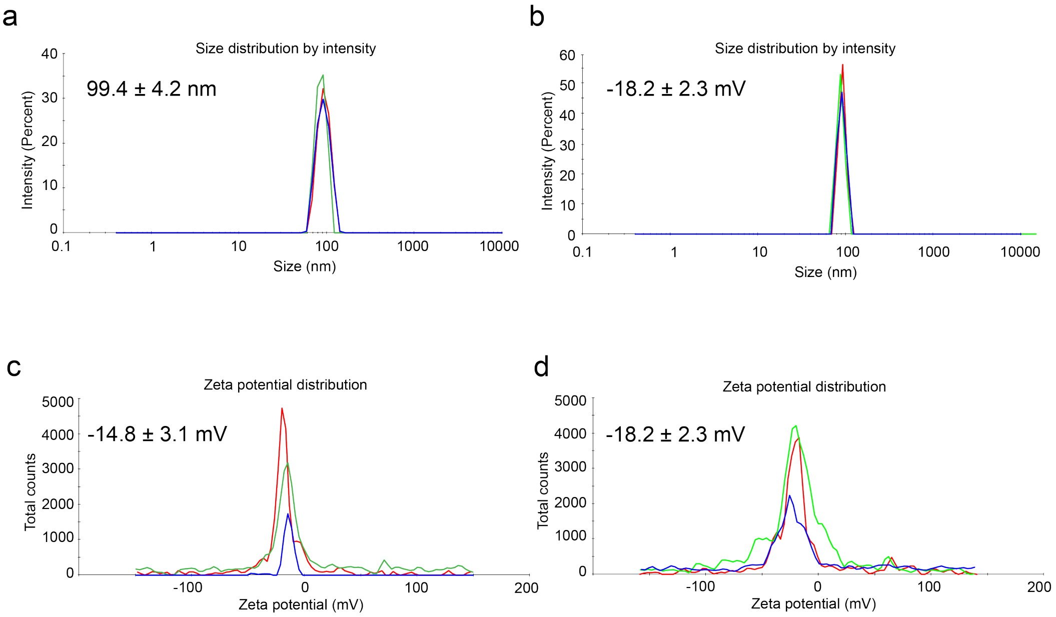


**Supplementary Fig. 5. Size and Zeta potential distribution of CFO-BTO and Alexa Fluor 594-labeled CFO-BTO in PBS. a,** Particle size distribution and (**c**) zeta potential measurements of CFO-BTO in PBS (n = 3). **b,** Particle size distribution and (**d**) zeta potential measurements of Alexa Fluor 594-labeled CFO-BTO in PBS (n = 3).

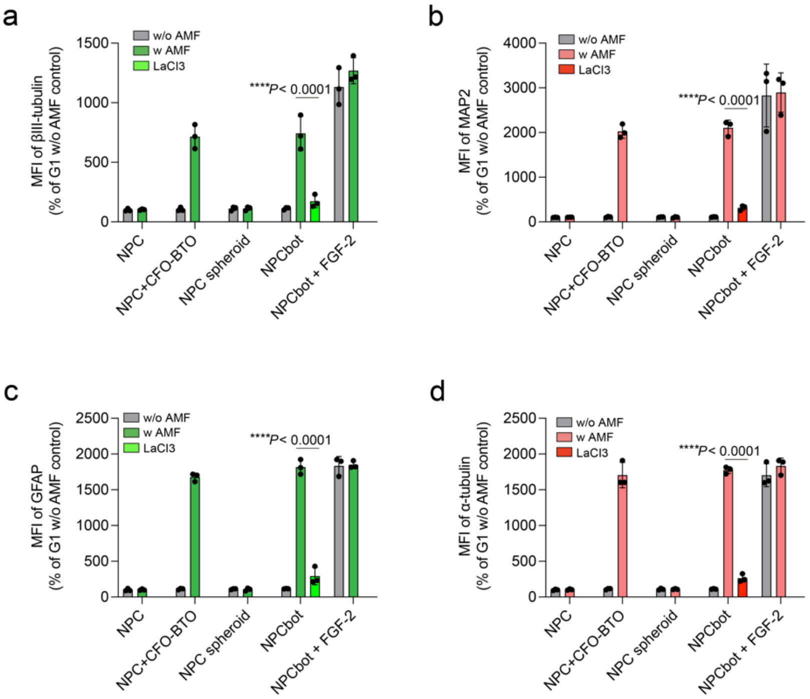


**Supplementary Fig. 6. Quantitative assessment through mean fluorescence intensity (MFI) across all samples based on the immunofluorescence results.** n = 3, means ± s.e.m. Statistical significance was calculated via one-way ANOVA with a Tukey post-hoc test. **** p< 0.0001 versus control.
